# Gene body H2B monoubiquitylation regulates gene-selective RNA Polymerase II pause release and is not rate limiting for transcription elongation

**DOI:** 10.1101/035386

**Authors:** Gilad Fuchs, Eran Rosenthal, Debora-Rosa Bublik, Tommy Kaplan, Moshe Oren

## Abstract

Histone H2B monoubiquitylation (H2Bub1) is localized preferentially to transcribed regions of genes and spreads concomitantly with the progression of RNA polymerase II (Pol II). In mammalian cells, H2Bub1 levels are highly correlated with transcription elongation rates, consistent with the general belief that H2Bub1 facilitates the elongation process. Yet, a causative role of H2Bub1 in regulating elongation rates within live cells remains to be proven. Using our recently developed 4sUDRB-seq method, we examined the impact of H2Bub1 downregulation, through silencing of its cognate E3 ubiquitin ligase RNF20, on genomewide transcription elongation rates. Surprisingly, H2Bub1 downregulation had no measurable effect on global elongation rates. Instead, it led to upregulation of over 1,000 genes by altering their Pol II pause release times; notably, those genes are characterized by the presence of H2Bub1 in relatively close proximity to the paused Pol II. Conversely, another set of genes was downregulated upon partial H2Bub1 depletion, and in those genes H2Bub1 appeared to be required for efficient recruitment of Pol II to the promoter region. Overall, our data shed new light on the molecular mechanisms by which H2Bub1 regulates gene expression and imply that the role of H2Bub1 in transcription elongation should be reconsidered.

**AUTHOR SUMMARY:** Transcription elongation is an important component of the gene expression process. Numerous factors and chromatin modifications, including some that are misregulated in various human diseases, have been suggested to regulate transcription elongation. New methods to measure genomewide transcription elongation rates now enable, for the first time, to determine how a specific factor affects transcription elongation and what is the outcome of its misregulation. Using such method, we examined the role of one specific chromatin modification, histone H2B monoubiquitination (H2Bub1), in regulating transcription. Strikingly, although H2Bub1 is widely believed to serve as a regulator of transcription elongation, its downregulation did not affect genomewide elongation rates. Instead, we found that H2Bub1 regulates the expression of distinct subsets of genes by either promoting recruitment of the transcription machinery or, conversely, favoring the pausing of this machinery shortly after initiation of transcription. Our findings demonstrate that the use of genomewide elongation rate measurements can redefine the true roles of putative transcription elongation factors. Furthermore, they provide a new understanding of the functions of H2Bub1 and its impact on gene expression patterns, which is of particular interest because H2Bub1 is often downregulated in human cancer.

## INTRODUCTION

Histones are widely modified post-translationally and have various roles in regulating molecular and cellular functions (1–6). One such modification is monoubiquitylation of histone H2B on lysine 123 in yeast or lysine 120 in mammals (H2BK120ub1, hereafter H2Bub1), carried out by the E3 ubiquitin ligase Bre1 in yeast (7, 8), and mainly by the heteromeric hBre1 (RNF20)/RNF40 complex in mammalian cells (9, 10). Additionally, H2B can be ubiquitylated on Lys34 (11).

H2Bub1 has been linked to many molecular processes such as transcription (12–18), RNA processing and export (19–22), DNA damage response (23–26), DNA replication (27) and nucleosome positioning (28). Given this multitude of processes, it is not surprising that misregulation of H2Bub1 was found to affect development, stem cell differentiation (29–31) and viral infection outcome (32, 33), and has been broadly implicated in cancer (34–37).

Among H2Bub1-regulated processes, the most extensively studied is transcription (36, 38, 39). The strong connection between H2Bub1 and transcription in mammals was demonstrated at various levels. First, H2Bub1 is most abundant within the transcribed regions of genes (40) and its local levels correlate with both extent of gene expression and transcription elongation rate (41). The tight link between H2Bub1 and RNA polymerase II (Pol II) elongation rate is also supported by its relatively low levels in exonic regions (41), where Pol II frequently pauses (42–45). Furthermore, real-time tracking of transcription elongation progress in long genes showed that H2Bub1 spreads concomitantly with Pol II movement (41). Of note, in *in vitro* transcription assays H2Bub1 was shown to regulate transcription cooperatively with the Facilitates chromatin transcription (FACT) and Polymerase Associated Factor (PAF) complexes, mainly at the elongation step, by enabling faster movement of Pol II through chromatin (14).

Intriguingly, *we* have previously observed that transient RNF20 knockdown in HeLa cells, which leads to a marked reduction in overall H2Bub1 levels, impacted the steady state levels of only a minority of transcripts (34). Two subgroups of genes were affected by the knockdown: one subgroup consisted of ~1000 genes downregulated upon RNF20 depletion (”RNF20-dependent”), while a similar number of genes were upregulated (“RNF20-suppressed”). In subsequent work, we found that H2Bub1 interferes with the recruitment to chromatin of the transcription elongation factor TFIIS; this provided a plausible explanation for the behavior of at least some of the RNF20-suppressed genes, which appear to require TFIIS for their efficient transcription (46).

Despite the reported links between H2Bub1 and transcription elongation, to date there is no direct *in vivo* evidence that H2Bub1 is functionally needed for transcription elongation in mammalians cells. Taking advantage of our recently developed 4sUDRB-seq method for genomewide measurement of transcription elongation rates (47, 48), we tested the direct effect of RNF20 knockdown and H2Bub1 downregulation on the elongation rates of thousands of genes. Surprisingly, even though we confirmed the effects of RNF20/H2Bub1 depletion on the expression of distinct subsets of genes, we did not observe a significant effect of such depletion on genomewide elongation rates, not even among the aforementioned groups of RNF20-regulated genes. We therefore explored alternative mechanisms that might explain the effect of RNF20/H2Bub1 depletion on gene expression. Interestingly, we identified a novel role of H2Bub1 in regulating promoter-proximal Pol II pausing in RNF20-suppressed genes. Conversely, in RNF20-dependent genes H2Bub1 appears to be required for efficient recruitment of Pol II to their promoter regions. Altogether, our findings question the importance of H2Bub1 for generic transcription elongation in mammalian cells and shed new light on the molecular mechanisms by which H2Bub1 regulates transcription.

## RESULTS

### Identification of H2Bub1-regulated genes by mRNA-seq and 4sU-seq in combination with RNF20 knockdown

Employing microarray technology, we previously reported that H2Bub1 depletion in HeLa cells affects the transcription of distinct subsets of genes, either positively or negatively (34). We now revisited these findings, using RNA-seq technology. HeLa cells were transfected with pools of RNF20-specific siRNA (siRNF20) oligos or control siRNA (siLacZ) oligos, and polyadenylated RNA was subjected to mRNA-seq analysis. As expected, RNF20 knockdown led to a substantial reduction of H2Bub1 (Fig. 1A). In agreement with our previous study (34), while the majority of genes showed no significant change in mRNA levels, 2343 genes were upregulated at least 1.3-fold and 1371 genes were downregulated at least 1.3-fold (Fig. 1B). The effects of RNF20 knockdown on the expression of representative genes were further validated by qRT-PCR (Fig. S1A).

**Figure 1.**
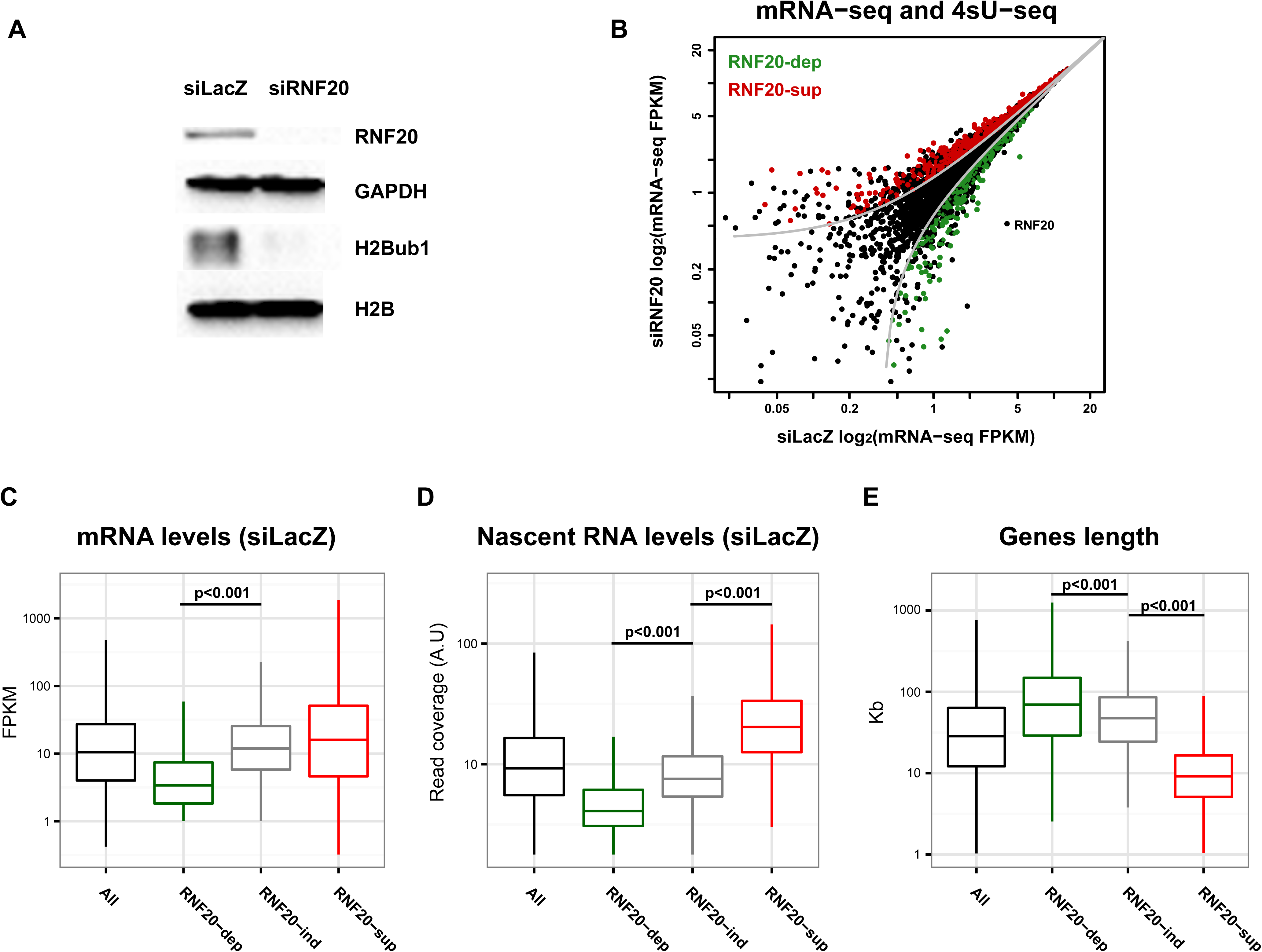
Identification and characterization of H2Bub1-regulated genes in HeLa cells. (A) HeLa cells were transfected with siRNA oligonucleotides (SMARTpool) directed against RNF20 (siRNF20) or against LacZ (siLacZ) as control, and harvested 48 hr later for Western blot analysis with the indicated antibodies. (B) Cells were transfected as in (A). 72 hours later, cells were labeled for 12 min with 1mM 4sU and harvested. For 4sU-seq, RNA was isolated, biotinylated *in vitro* and enriched on streptavidin beads. For mRNA-seq, total polyadenylated (polyA+) RNA (non 4sU-enriched) was selected with oligo-dT magnetic beads. Both polyA+ and 4sU-enriched RNA were next subjected to high throughput sequencing. Shown is a scatterplot of the mRNA-seq analysis of the siLacZ and siRNF20 samples. Grey lines mark a fold change threshold of 1.3. Genes that were also significantly upregulated or downregulated in the 4sU-seq analysis are marked in red or green, respectively. Axes are in log scale. (C) Box plot showing mRNA-seq FPKM levels in control (siLacZ-transfected) HeLa cells of all annotated RefSeq genes (‘All’), downregulated (‘RNF20-dependent’) genes, unchanged (‘RNF20-independent’) genes and upregulated genes (‘RNF20-suppressed’). PolyA+ mRNA levels of RNF20-dependent genes are significantly lower than those of RNF20-independent genes (Kolmogorov–Smirnov test, *P* < 2.2e-16). The Y-axis is in log scale. Outliers are excluded (see methods). (D) Same as (C) but for 4sU-seq. RNF20-suppressed genes have significantly higher nascent RNA levels than RNF20-independent genes (Kolmogorov-Smirnov test, *P* < 2.2e-16), while RNF20-dependent genes have significantly lower nascent RNA leves (Kolmogorov–Smirnov test, *P* < 2.2e-16). (E) Same as (C) but for gene lengths. RNF20-dependent genes are significantly longer than RNF20-independent genes (Kolmogorov–Smirnov test, *P =* 9.8e-13), while RNF20-suppressed genes are significantly shorter (Kolmogorov–Smirnov test, *P* < 2.2e-16).

Steady state mRNA levels represent a balance between transcription and degradation rates (49–51) and do not necessarily reflect the actual rates of active transcription per gene. To determine directly the effects of H2Bub1 downregulation on genomewide transcription rates, we therefore measured nascent RNA levels. As nascent RNA constitutes only a small portion of the total RNA within cells, we enriched the nascent RNA by treating the cells with a short pulse of 4-thiouridine (4sU), which is incorporated into newly transcribed RNA (52, 53). Nascent transcripts were then enriched by *in vitro* biotinylation, purified on streptavidin beads, and sequenced (4sU-seq). By integrating mRNA-seq and 4sU-seq data in RNF20-depleted versus control cells, we were able to identify genes whose transcription is indeed regulated by H2Bub1 (Fig. 1B, Fig. S1B and Table S1). Out of the 2343 genes whose steady state mRNA levels were upregulated upon RNF20 knockdown, 1344 also showed elevated transcription (Fig.1B, red), whereas out of the 1371 genes with downregulated mRNA, reduced transcription could be confirmed for only 432 (Fig.1B, green). For subsequent analyses, only genes for which both mRNA and nascent RNA levels were either up- or downregulated upon RNF20 knockdown were considered “RNF20-suppressed” and “RNF20-dependent”, respectively. These combined gene lists correlate significantly with those identified in our previous study (34) (Hypergeometric *p*-value = 0 and *P ≤* 6.6e-11, respectively).

We next searched for distinctive properties of each group of RNF20-regulated genes. To that end, we compared the basal levels of mRNA and nascent RNA between the two groups. As controls, we chose a third group of genes, similar in size, whose mRNA and nascent RNA levels were unchanged following RNF20 depletion (“RNF20-independent”, n = 978, Fig. S1B). Interestingly, while the average steady state mRNA levels of RNF20-suppressed genes are relatively similar to global mRNA levels (“All”) and to those of RNF20-independent genes (Fig. 1C), these RNF20-suppressed genes exhibit significantly higher nascent transcript levels, ~3 fold higher on average than RNF20-independent genes (Kolmogorov–Smirnov test, *P ≤* 2.2e-16), suggesting that their mRNAs may have relatively short half-lives or, alternatively, may have slower rates of splicing or degradation of introns. On the other hand, the RNF20-dependent genes are characterized by both lower steady state mRNA and lower nascent RNA levels, relative to RNF20-independent genes.

During differentiation of human embryonal carcinoma stem cells, transcriptional induction of long genes is selectively sensitive to RNF20 knockdown (30). In agreement, while the median length of all human genes is ~16Kb, RNF20-dependent genes in HeLa cells tended to be significantly longer (median length of 70Kb, Kolmogorov-Smirnov test, *P ≤* 1e-12, Fig. 1D). Notably, RNF20-suppressed genes were significantly shorter (median length of 9Kb, Kolmogorov–Smirnov test, *P ≤* 2.2e-16, Fig. 1D). As short genes tend to be highly transcribed (54), the shorter size of the RNF20-suppressed genes is in agreement with their relatively high transcription rates (Fig. 1C).

Interestingly, pathway enrichment analysis indicated that many ribosomal proteins and translation-related functions are enriched in the RNF20-suppressed group (Fig. S1C). In contrast, the RNF20-dependent genes did not show strong enrichment for a single specific pathway (Fig. S1C).

### H2Bub1 downregulation does not affect genomewide transcription elongation rates

H2Bub1 stimulates transcript elongation *in vitro* (14) and its abundance on specific genes in human cells correlates significantly with elongation rates (41). Hence, it seemed plausible that the effects of H2Bub1 depletion on the two groups of RNF20-regulated genes might be explained through changes in their transcription elongation rates. However, there is so far no direct evidence that H2Bub1 regulates transcription elongation in intact cells. To search for such evidence, we took advantage of the 4sUDRB-seq method, which measures genomewide transcription elongation rates (47, 48). Specifically, transcription was inhibited by DRB in control and RNF20-depleted cells, and nascent RNA was quantitatively profiled by 4sU-sequencing 8 and 12 minutes after removal of DRB (to reinitiate transcription). Surprisingly, despite a substantial decrease in *RNF20* mRNA (Fig. S2A), H2Bub1 downregulation did not significantly affect global transcription elongation rates (Figure 2A and Fig. S2B). Likewise, no significant effects were observed when elongation rates of RNF20-dependent or RNF20-suppressed genes were examined separately (Fig. 2B). The conclusions from 4sUDRB-seq were further supported by qRT-PCR-based elongation rate measurements of specific genes after DRB removal (Fig. S2C).

**Figure 2.**
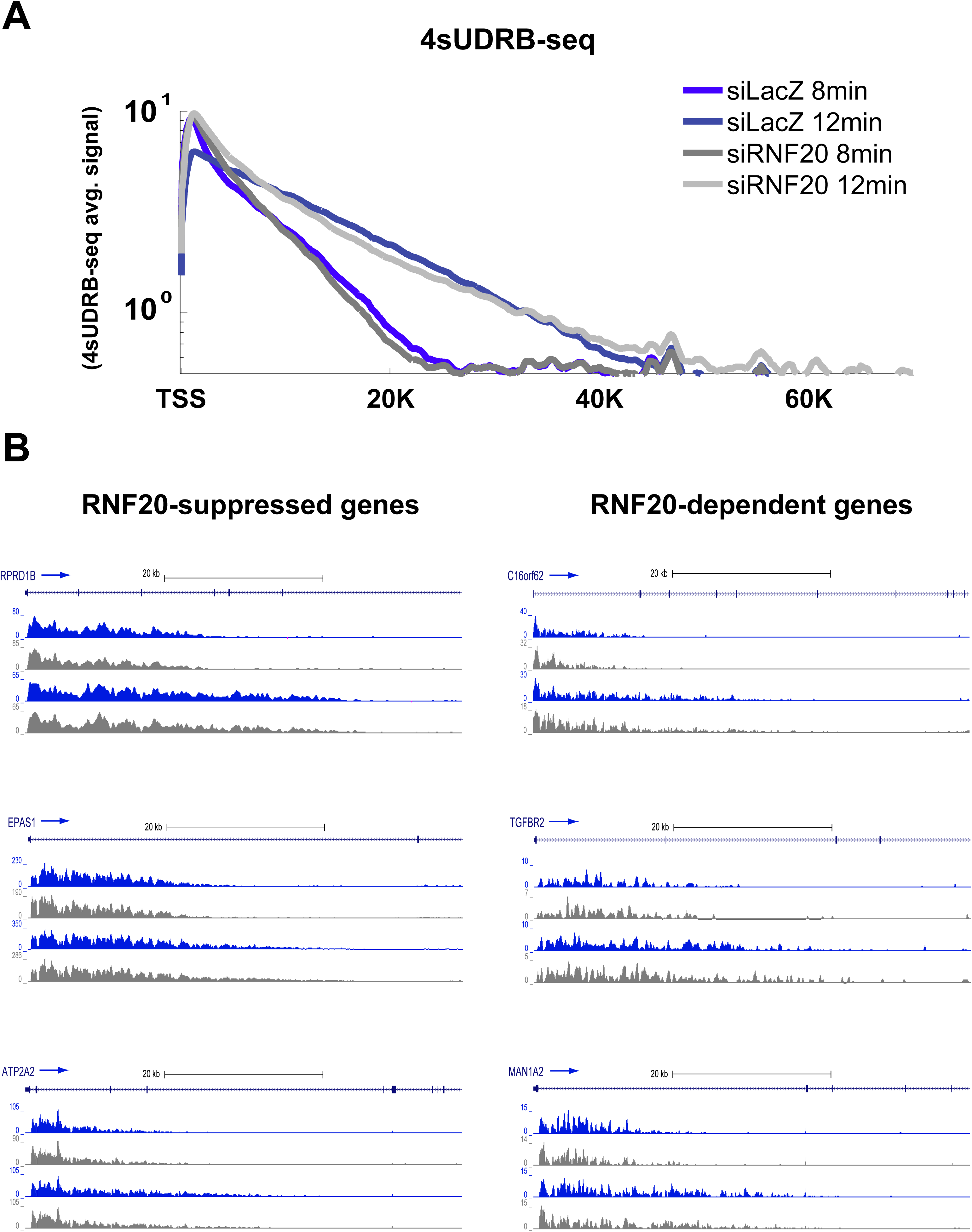
H2Bub1 depletion does not affect genomewide transcription elongation rates. (A) Average 4sUDRB-seq signals as a function of distance from the TSS in RNF20-depleted (siRNF20) or control (siLacZ) HeLa cells treated for 3 h with 100 mM DRB and harvested 8 or 12 min after DRB removal, Only genes longer than 35 Kb (n=9532) were included. (B) 4sUDRB-seq reads distribution along representative RNF20-suppressed and RNF20-dependent genes. Shown is 4sUDRB-seq data from cells harvested 8 (top) or 12 (bottom) minutes following DRB release, for siRNF20 (grey) or control (siLacZ, blue) cells. Exonic 4sU-seq reads were excluded from the analysis, to minimize contamination from mature RNAs. Arrows mark the direction of transcription.

Together, these results suggested that the observed changes in transcription rates following RNF20 depletion are probably mediated through a mechanism different than transcription elongation.

### H2Bub1 regulates promoter-proximal pausing of RNF20-suppressed genes

The transcription process typically involves recruitment of Pol II, initiation and gene entry, promoter-proximal pausing and release, productive elongation and termination (55). Given our finding that H2Bub1 depletion does not affect productive elongation, we sought to examine whether other steps in the transcription process are regulated by H2Bub1 in a manner that might explain the observed effects on transcription and gene expression. To that end, we employed Chromatin Immunoprecipitation coupled with high throughput sequencing (ChIP-seq) to analyze the genomewide distribution of Pol II following RNF20 knockdown. N-terminal-specific antibodies, which bind the large subunit of Pol II regardless of its phosphorylation state, were used for that purpose. We obtained over 180 million 50bp single-end sequence reads (Illumina HiSeq 2500) and mapped them to the genome (UCSC hg19 release) using BOWTIE (56) resulting in 150 million uniquely mapped reads. Analysis of the data revealed a significant selective reduction in Pol II abundance near the transcription start site (TSS) of the RNF20-dependent genes upon RNF20 knockdown (Fig. 3A, Kolmogorov–Smirnov test, *P ≤* 2.2e-16). Consistent with the reduction in the TSS region, Pol II levels decreased similarly also in the bodies of those genes (Fig. 3B). The decrease in Pol II in both TSS and gene body regions suggests that in RNF20-dependent genes, H2Bub1 is mainly needed for Pol II recruitment.

**Figure 3.**
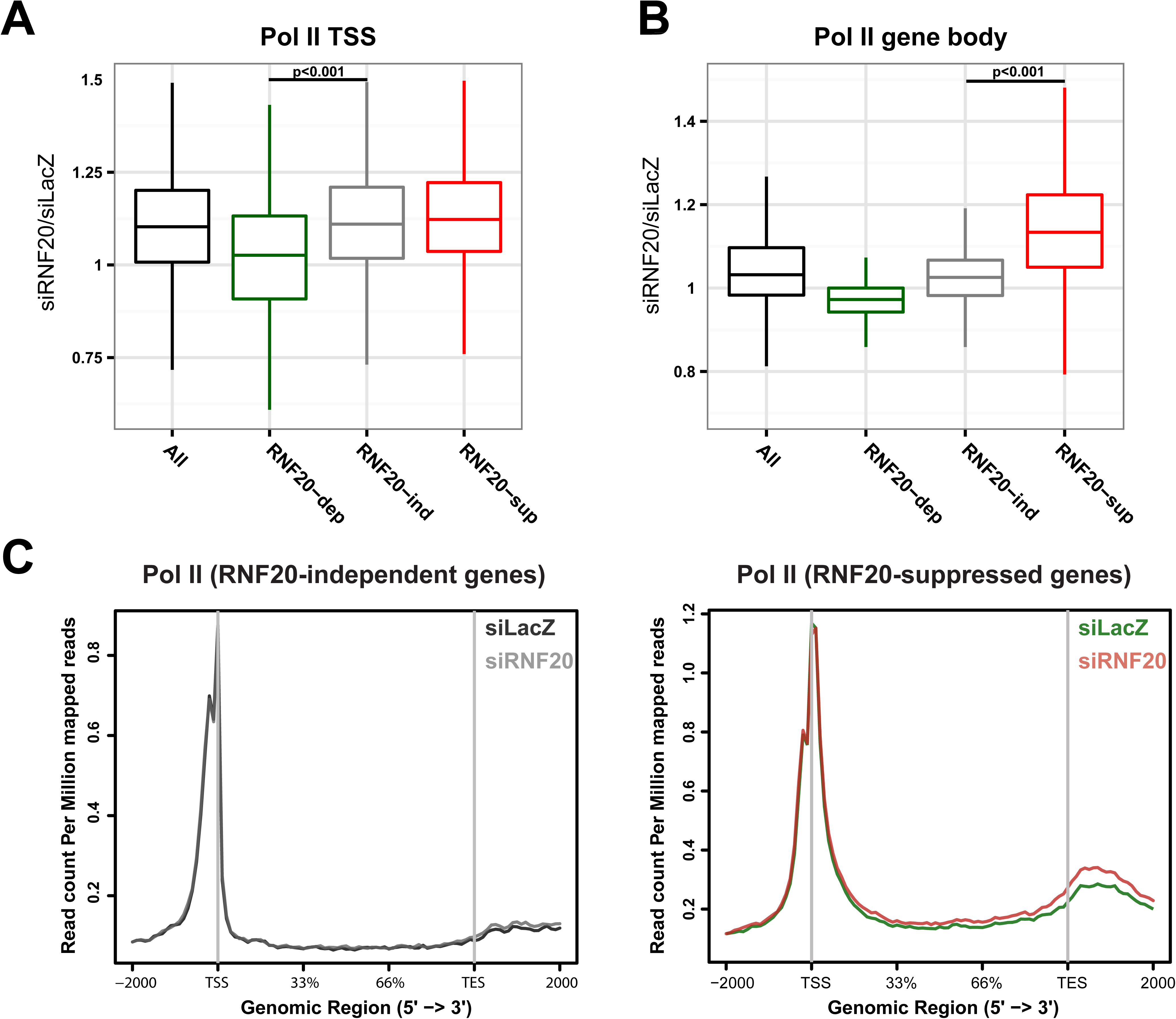
RNF20 regulates Pol II recruitment to RNF20-dependent genes and promoter proximal pausing within RNF20-suppressed genes. (A) Box plot of the ratio between Pol II levels (from ChIP-seq) in siRNF20 cells versus control (siLacZ) cells in the region near the TSS (+/− 500 bps) of all annotated RefSeq genes with significant signal (‘All’), as well as in RNF20-dependent, RNF20-independent and RNF20–suppressed genes. ChIP was performed on HeLa cells 72 hr following siRNA transfection. Pol II levels are significantly reduced near the TSS of RNF20-dependent genes following RNF20 depletion (Kolmogorov–Smirnov test, *P* < 2.2e-16) relative to RNF20-independent genes. Outliers are excluded (see methods). (B) Same as (A) but gene body Pol II levels were analyzed. Gene body Pol II levels are significantly elevated in RNF20-suppressed genes following RNF20 depletion (Kolmogorov–Smirnov test, *P* < 2.2e-16). (C) Normalized Pol II ChIP-seq read densities across the TSS, gene bodies and transcription end sites (TES) in RNF20-suppressed (left) and RNF20-independent (right) genes. Gene bodies were normalized to 0–100% as relative position.

Intriguingly, although the transcription rates of RNF20-suppressed genes increased significantly upon RNF20 knockdown, these genes showed only a very mild increase in the recruitment of Pol II to the TSS region, relative to the genomewide trend (Fig. 3A). In contrast, Pol II abundance increased significantly along their gene bodies (Fig. 3B, Kolmogorov–Smirnov test, *P* ≤ 2.2e-16). This suggests that in RNF20-suppressed genes, H2Bub1 regulates transcription by specifically inhibiting the transition from a paused polymerase into a state of productive elongation. A similar trend was also revealed by a metagene analysis of Pol II average occupancy: while RNF20-independent genes showed almost no change in Pol II distribution upon RNF20 knockdown (Fig. 3C, left panel), RNF20-suppressed genes (right) showed a mild progressive increase in Pol II levels along the gene body, becoming more obvious near the transcription end site (TES).

In sum, our findings suggest that H2Bub1 restricts the transcription of RNF20-suppressed genes by inhibiting promoter-proximal Pol II pause release.

### H2Bub1 is localized in closer proximity to paused Pol II in RNF20-suppressed genes

As H2Bub1 downregulation affected Pol II pause release in only a subset of genes, we looked for distinctive features of the RNF20-suppressed genes that might explain why they are selectively affected. A plausible answer seemed to be suggested by a recent H2Bub1 interactome screen, which identified DRB sensitivity-inducing factor (DSIF) and negative elongation factor (NELF) as possible H2Bub1 interactors (58). As both DSIF and NELF have pivotal roles in regulating Pol II pause release (59, 60), we compared their distribution (data from (61)) in RNF20-suppressed and RNF20-independent genes, stratified for similar transcription levels (n = 810 and n = 507, respectively). However, we found no significant differences between the two groups of genes with respect to the distribution or extent of binding of DSIF and NELF (Fig. S3A).

Intriguingly, interrogation of published HeLa H2Bub1 ChIP-seq data (62) revealed that gene body H2Bub1 was positioned in closer proximity to the TSS of RNF20-suppressed genes as compared to RNF20-independent (Fig. 4A) or dependent (Fig. S3B) genes. This is not due to differences in overall H2Bub1 occupancies, which became comparable at about +1.5Kb into the gene bodies (Fig. S3C). Moreover, the differences in 5’ H2Bub1 localization could not be explained by different positions of the first nucleosome (Fig. S3D, data from (63)). H2Bub1 is relatively low in exons and enriched in the first intron (41), raising the possibility that the RNF20-suppressed genes might be characterized by short first exons and thus a shorter distance between the TSS and the first exon-intron boundary. Indeed, this was found to be the case (Fig.4B, Kolmogorov–Smirnov test, *P* = 3.6e-10 and *P* = 9.7e-6).

**Figure 4.**
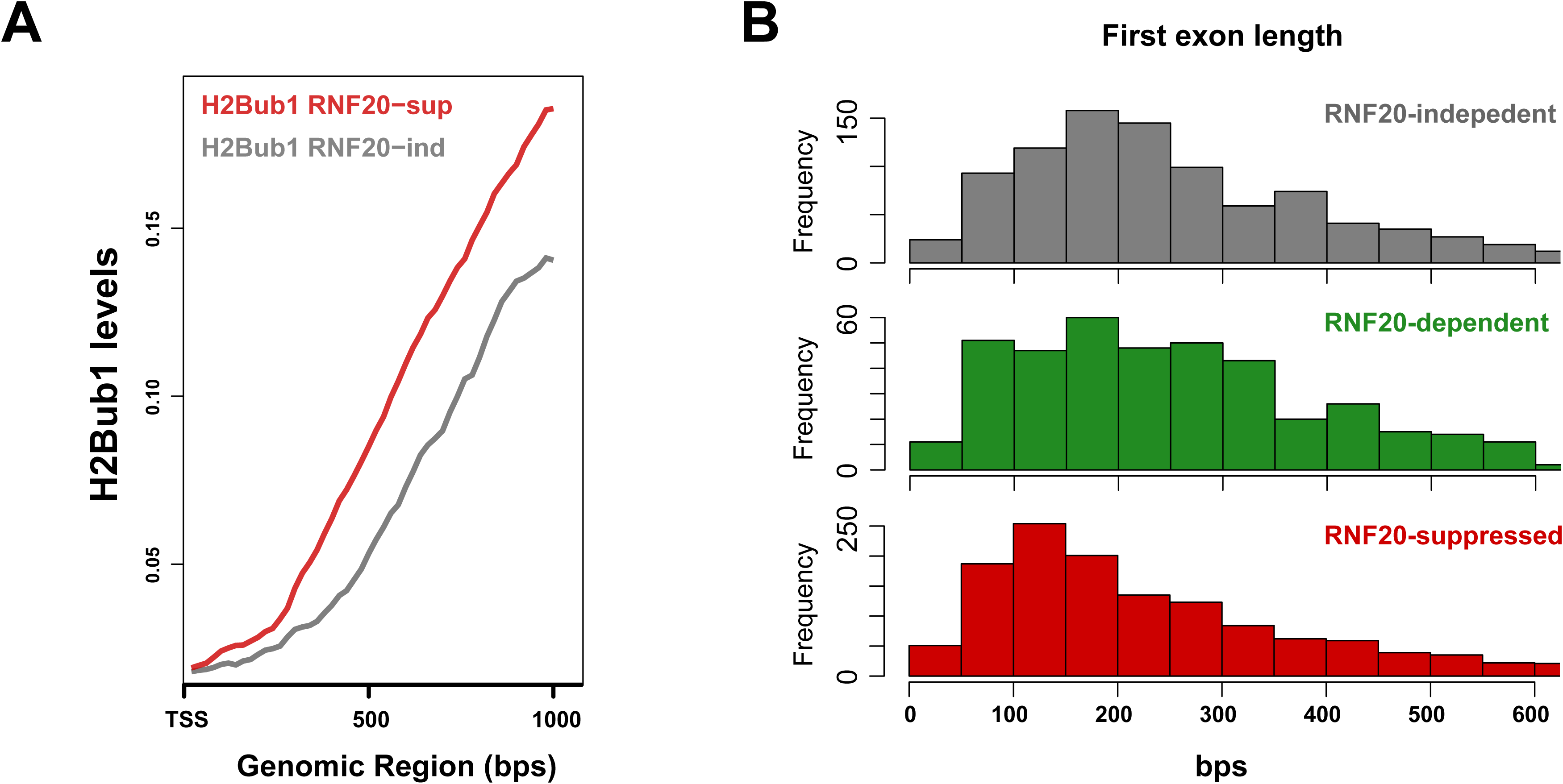
H2Bub1 is localized in closer proximity to paused Pol II in RNF20-suppressed genes. (A) Comparison of the distributions of H2Bub1 (data from (62)) read densities near the TSS of RNF20-suppressed and RNF20-independent genes. Genes with relatively similar transcription levels from both groups (n = 810 and n = 507, respectively) were selected for this analysis. (B) First exon length distributions of RNF20-independent, -dependent and -suppressed genes. Exons longer than 600 bps were excluded from the analysis. First exons of RNF20-suppressed genes are significantly shorter than of RNF20-independent and RNF20-dependent genes (Kolmogorov-Smirnov test, *P* = 3.6e-10 and *P* = 9.7e-6).

Together, these observations suggest that the proximity of the first H2Bub1 to the TSS might contribute to Pol II pausing on the RNF20-suppressed genes, either directly by steric hindrance or indirectly by affecting the positioning of other associated proteins.

## DISCUSSION

Cell-free transcription experiments have provided compelling evidence that H2Bub1 is required for optimal transcription elongation (14). However, by combining direct measurement of genomewide elongation rates, mRNA-seq and Pol II distribution data, we found surprisingly that despite its tight coupling to transcription elongation, H2Bub1 is not rate limiting for transcription elongation in human cells (Fig. 2). However, we uncovered new roles of H2Bub1 in Pol II recruitment and regulation of pause release in a gene-specific manner. While the increase in Pol II in gene bodies of RNF20-suppressed genes (Fig. 3B) might have also been interpreted as slower elongation, 4sUDRB-seq excluded this possibility.

Interestingly, in RNF20-dependent genes RNF20 knockdown caused a decrease in Pol II levels both near the TSS and within the gene body (Fig. 3A). Such decrease might potentially be explained by either reduced recruitment of Pol II to the gene or a shorter retention time of Pol II in paused state (64). As H2Bub1 may be a prerequisite for H3K4 trimethylation (H3K4me3) in at least some genes (65, 66), and H3K4me3 regulates preinitiation complex assembly through TFIID (67), it is possible that H3K4me3 plays a role in mediating the effect of H2Bub1 in RNF20-dependent genes. Moreover, RNF20-dependent genes rely on the SWI/SNF chromatin remodeling complex for their efficient transcription, and H2Bub1 is needed for the optimal recruitment of SWI/SNF to those genes (58). Hence, H2Bub1-mediated recruitment of SWI/SNF to the TSS regions may serve to adjust the 5’ nucleosome landscape of RNF20-dependent genes in order to allow efficient recruitment of Pol II. It should also be noted that we cannot presently exclude formally the possibility that other unknown targets of RNF20, different from H2B, also contribute to some of the observed effects of RNF20 knockdown.

As regards the RNF20-suppressed genes, it has long been known that RNF20 knockdown, with subsequent H2Bub1 downregulation, enables the more efficient transcription of those genes (34). However, the exact stage of the transcription process at which this effect was exerted remained unclear, leaving transcription elongation as the most likely candidate. Unexpectedly, the present study indicates that RNF20, and thus presumably H2Bub1, actually exerts its effect on those genes by enhancing promoter-proximal Pol II pausing. The question still remains how the gene-specific nature of this effect is determined. We believe that an important clue is provided by the observation that, in the RNF20-suppressed genes, H2Bub1 is localized in relatively close proximity to the TSS. This distinctive feature may be dictated by the fact that RNF20-suppressed genes possess characteristic short first exons (Fig. 4B); as reported previously, promoter-proximal accumulation of H2Bub1 starts at the boundary between the first exon and the first intron (41). An alternative, and not mutually exclusive explanation may be provided by the recent study of Rhee et al. (68) who found that yeast H2Bub1 is distributed asymmetrically within the first nucleosome. It is possible that such asymmetry also exists in mammalian chromatin, but only in a subset of genes, and in the RNF20-suppressed genes the H2B monomer located closer to the TSS in the first nucleosome is preferentially ubiquitinated. However, due to the large size of the human genome and the limited resolution of our sonication-based data, we can not presently determine whether this is indeed the case.

How can the proximity of H2Bub1 to the TSS favor Pol II pausing? Ubiquitin is a bulky modification, and thus its proximity to Pol II may physically inhibits the release of Pol II from its paused state into productive elongation. Alternatively, H2Bub1 may regulate Pol II pausing through affecting the localized recruitment of additional factors that either promote pausing or are required for efficient pause release. In particular, we previously found that TFIIS is needed for efficient transcription of RNF20-suppressed genes and H2Bub1 prevent TFIIS recruitment to chromatin (46). As TFIIS was reported to regulate Pol II pause release (69), it is possible that close proximity of H2Bub1 to the TSS physically interferes with TFIIS recruitment and thereby slows down the release of Pol II from its paused state. Furthermore, as H2Bub1 was shown in yeast to prevent the binding of Ctk1 and phosphorylation of Pol II on Ser2 (40), it is possible that H2Bub1 also decreases the recruitment of the mammalian positive transcription elongation factor (P-TEFb) to RNF20-suppressed genes, resulting in increased pausing.

Although our findings implicate H2Bub1 as gene-specific regulator of transcription initiation and promoter-proximal pause release, the role of H2Bub1 within the gene bodies, where it is actually most abundant, remains to be figured out. Clearly, H2Bub1 does not simply regulate elongation rates; hence, it may have a different, yet to be determined role in later stages of the transcription process. One possible role might be suggested by the recently identified importance of the Polymerase-Associated Factor (PAF) complex, known to promote H2Bub1 (70), towards protecting fission yeast genes from small RNA-mediated epigenetic silencing (71). Moreover, we believe that the present findings should encourage others to revisit the roles of other proposed elongation factors by testing directly how their depletion affects transcription elongation rates.

## MATERIALS AND METHODS

### Cell culture and treatments

Human cervical carcinoma HeLa cells were grown in DMEM with 10% bovine serum supplemented with antibiotics and maintained at 37°C. 5,6-dichlorobenzimidazole 1-β-d-ribofuranoside (DRB) was purchased from Sigma (D1916), and used at a final concentration of 100μM. 4-thiouridine (4sU) was purchased from Sigma (T4509) and used at a final concentration of 1mM. SMARTpool oligonucleotides for siRNA transfection were purchased from Dharmacon. Transfections were carried out with Dharmafect 1 reagent according to the manufacturer’s protocol. All siRNA oligos were used at a final concentration of 25 nM.

### RNA purification and quantitative RT-PCR

For quantitative reverse transcription (RT)-PCR analysis, total RNA was extracted with the miRNeasy kit’s procedure (Qiagen). Two micrograms of each RNA sample was reverse transcribed with Moloney murine leukemia virus reverse transcriptase (Promega) and random hexamer primers (Applied Biosystems). Real-time PCR was done in a StepOne real-time PCR instrument (Applied Biosystems) with SYBR Green PCR supermix (Invitrogen). Primers used in this study are listed in Table S2.

### Antibodies

The following commercial antibodies were used: anti-H2B (07-371, Millipore), anti-GAPDH (MAB374, Millipore), anti-Pol II (SC-899, Santa Cruz). Anti-H2Bub1 is described in Minsky et al. (72).

### ChIP and high-throughput sequencing

Cells were fixed by adding one-tenth volume of an 11% formaldehyde solution directly to the medium for 10 min at room temperature, washed twice and harvested. To extract chromatin, cells were resuspended in 1 mL cell lysis buffer (50 mM PIPES, 85 mM KCl, 0.5% NP-40); nuclei were pelleted by centrifugation and resuspended in 100 μL of nuclei lysis buffer (50 mM Tris-HCl, 10 mM EDTA, 1% SDS). Next, samples were diluted 1:4.5 in dilution buffer (0.01% SDS, 1.1% Triton X-100, 1.2 mM EDTA, 16.7 mM Tris-HCl, 167 mM NaCl) and subjected to 30 cycles of sonication; each cycle included 30 sec of sonication and a 30 sec interval. Sonicated chromatin was incubated overnight at 4°C with a 50% slurry of protein A–Sepharose beads blocked with yeast transfer RNA (Sigma, R8759) together with the appropriate antibody. Immunoprecipitates were washed with dilution buffer and eluted with 50 mM NaHCO3, 1% SDS. DNase-free RNase and Proteinase K were added to the immunoprecipitated chromatin to final concentrations of 50 ng/mL and 200 ng/mL, respectively, followed by incubation for 3 h at 45°C. To complete reversal of crosslinks, samples were incubated overnight at 65°C. DNA was purified using the Qiagen PCR purification kit according to the manufacturer’s protocol. High-throughput sequencing was performed on 5ng DNA from two technical repeats of two independent experiments of Pol II ChIP, processed as previously described (73). Libraries were evaluated by Qubit and TapeStation. Sequencing libraries were constructed with barcodes to allow multiplexing of 8 samples on one lane. 20 million single-end 50-bp reads per sample were obtained in an Illumina HiSeq 2500 sequencer

### 4sUDRB-seq and mRNA-seq

Cells were treated as described in (48) except that the 4sU pulse was for 12 min. Biotinylation and purification of 4sU-labeled RNA from two independent experiments was done as described in (48). cDNA libraries were prepared with Illumina TruSeq RNA sample preparation v2 kit according to the manufacturer’s instructions but without the polyA isolation stage. Of note, strand specificity is not achieved with this kit. Random hexamers were used for reverse transcription. Libraries were pooled and sequenced on an Illumina HiSeq 2500 system with single-end 100bp reads. For mRNA-seq, total non-biotinylated RNA from two independent experiments was used to generate cDNA libraries using the Illumina TruSeq RNA sample preparation v2 kit according to the manufacturer’s instructions. Radom hexamers were used for reverse transcription. Libraries were pooled and sequenced on an Illumina HisSeq 2500 system with paired-end 50bp reads. Sequenced reads were mapped to the genome (UCSC release hg19) using TopHat (74), with an expected paired-end inner distance of 200, and gene expression was estimated using Cufflinks (75), with a single MLE iteration.

### Data analysis

In all boxplot figures outliers were defined by calculating 1.5 times the interquartile range (third quartile minus first quartile), creating limits by subtracting this value from the first quartile, and adding it to the third quartile.

### Elongation rates

For each gene, a discrete Hidden Markov Model of transcribed and non-transcribed states was used to define the transcribed regions, based on the 4sUDRB-seq data. The non-transcribed state is constrained to do self-transitions only. Each value is mapped to a discrete value based on thresholds of 5%, 50%, 75%, 90% percentiles of the data. The HMM is trained based on data of 8 minutes and 12 minutes, excluding exons and subtracting data of time = 0, to avoid reads where the DRB is less effective.

## ACKNOWLEDGMENTS

We thank Dror Hollander, Efrat Shema, Shlomit Gilad and Yoav Voichek for helpful discussions.

## SUPPORTING INFORMATION

**Figure S1. Characterization of RNF20-regulated genes. Related to Figure 1**

(A) qRT-PCR analysis of relative mRNA levels of representative RNF20-independent, - dependent, and –suppressed genes, 72 hr after transient transfection of HeLa cells with RNF20 siRNA (siRNF20) or LacZ siRNA (siLacZ) as control. Transcript levels in the siLacZ samples were set as 1. Values were normalized to 18S ribosomal RNA in the same sample. Bars indicate averages of values from duplicate qPCR reactions; error bars represent standard deviation. Similar data were obtained in four independent experiments. (B) Cells transfected as in (A) were labeled for 12 min with 1mM 4sU. For 4sU-seq, RNA was isolated, biotinylated, and enriched on streptavidin beads. For mRNA-seq, total polyadenylated (polyA+) RNA (non 4sU-enriched) was selected with oligo-dT magnetic beads. Both PolyA+ and 4sU-enriched RNA were next subjected to high throughput sequencing. Shown is a scatterplot of the mRNA-seq analysis of the siLacZ and siRNF20 samples. Grey lines mark a fold change threshold of 1.3. Genes that were also significantly upregulated or downregulated in the 4sU-seq analysis are marked in red or green, respectively. Genes with a siRNF20/siLacZ ratio lower than 1.1 but higher than 0.9 in both mRNA-seq and 4sU-seq were defined as RNF20-independent genes and marked in grey. (C) Pathway enrichment (ConsensusPathDB) of RNF20-suppressed and RNF20-dependent genes. Only pathways with FDR < 0.1 are shown.

**Figure S2. H2Bub1 depletion does not affect transcription elongation rates of RNF20-regulated genes. Related to Figure 2**

(A) Quantification of *RNF20* mRNA 72 hr after transient transfection of HeLa cells with *RNF20* siRNA (siRNF20) or LacZ siRNA (siLacZ) as control, after 3 h DRB treatment (DRB) and at the indicated time points after DRB removal. NT = non-treated. Values were normalized to *GAPDH* mRNA in the same sample. Bars indicate averages of values from duplicate qPCR reactions; error bars represent standard deviation. (B) Box plot showing the difference in elongation rates between siLacZ and siRNF20-transfected cells in all annotated RefSeq genes (‘All’), downregulated (‘RNF20-dependent’), unchanged (‘RNF20-independent’) and upregulated (‘RNF20-suppressed’) genes longer than 35 Kb. Outliers were excluded (see methods). (C) Cells were transfected as (A), and *TTC17* and *CADM1* pre-RNA levels at the indicated time points after DRB removal were quantified by qRT-PCR, employing intron-derived primers. All values were normalized to 16S ribosomal RNA in the same sample. Similar data were obtained in two independent experiments.

**Figure S3. Distribution of DSIF, NELF and nucleosomes in RNF20-regulated genes. Related to Figure 4**

(A) Distribution of DSIF and NELF (data from (61)) ChIP-seq read densities around the TSS of RNF20-suppressed and RNF–independent genes, pre-selected for similar expression levels (n = 810 and n = 507, respectively). (B) Distribution of H2Bub1 ChIP-seq read densities around the TSS of RNF20-suppressed and RNF20-dependent genes. (C) As in (B) but comparing RNF20-suppressed and RNF20-independent genes and pre-selected for similar expression levels. (D) MNase-seq (data from (63)) read densities around the TSS of RNF20-suppressed and RNF20-independent genes. The first nucleosome is annotated as +1.

**Table S1. List of RNF20-suppressed and RNF20-dependent genes**. Fold change of mRNA-seq and 4sU-seq is indicated as the ratio between siRNF20 and siLacZ.

**Table S2. List of primers used in this study.**

## DATA REPORTING

The sequencing data from this study have been submitted to the NCBI Gene Expression Omnibus (GEO; http://www.ncbi.nlm.nih.gov/geo/) under accession number GSE69738.

